# A flow extension tethered particle motion assay for single-molecule proteolysis

**DOI:** 10.1101/528919

**Authors:** Andrew A. Drabek, Joseph J. Loparo, Stephen C. Blacklow

## Abstract

Regulated proteolysis of signaling proteins under mechanical tension enables cells to communicate with their environment in a variety of developmental and physiologic contexts. The role of force in inducing proteolytic sensitivity has been explored using magnetic tweezers at the single-molecule level with bead-tethered assays, but such efforts have been limited by challenges in ensuring that beads are not restrained by multiple tethers. Here, we describe a multiplexed assay for single-molecule proteolysis that overcomes the multiple-tether problem using a flow extension (FLEX) strategy on a microscope equipped with magnetic tweezers. Particle tracking and computational sorting of flow-induced displacements allows assignment of tethered substrates into singly-captured and multiply-tethered bins, with the fraction of fully mobile, single-tethered substrates depending inversely on the concentration of substrate loaded on the coverslip. Computational exclusion of multiply-tethered beads enables robust assessment of on-target proteolysis by the highly specific tobacco etch virus protease and the more promiscuous metalloprotease ADAM17. This method should be generally applicable to a wide range of proteases and readily extensible to robust evaluation of proteolytic sensitivity as a function of applied magnetic force.

## Introduction

Force-dependent proteolysis of tension-sensing domains is a fundamental mechanism for signal transduction in biology ^1–4^. For example, in response to shear forces in the vasculature, von Willebrand Factor undergoes proteolysis in its force-sensing domain to regulate blood clotting ^5^. At sites of cell-cell contact, a mechanosensing domain in the Notch receptor undergoes regulated proteolysis in response to tension applied by bound ligand, influencing cell fate decisions during development ^6–7^. In the cellular response to the extracellular matrix, force-induced cleavage of talin is essential for mechanosensation and the adhesion response ^8^.

Given the importance of force-dependent proteolysis in biological signaling, we sought to develop a robust, single-molecule assay for proteolytic cleavage that can be readily adapted for probing proteolytic sensitivity in response to force. In this regard, single-molecule magnetic tweezers methods enable direct, multiplexed observations of tension-dependent biochemical processes, in cell-based assays or in purified systems *in vitro* ^9^. These approaches typically rely on the capture of a substrate between a probe and anchor surface, normally a colloidal bead and glass coverslip, respectively. One problem that can confound these assays is the simultaneous capture of one bead by more than one immobilized substrate molecule, the so called “multiple-tether problem,” which has not been directly accounted for in previous multiplexed single-molecule proteolysis studies ^6, 10^.

To address the multiple tethering problem, Dekker and coworkers used micropatterning to separate anchor points based on the contour length of a simple DNA substrate. Although the approach enriches for single tethers, it does not eliminate multiply tethered particles ^11^. In studying unzipping of DNA nanoswitches, Wong and colleagues used tethered particle motion and a length change signature to perform sorting for centrifugal force spectroscopy ^12^. This approach is highly specialized, however, and requires complex custom instrumentation that is not yet commercially available.

We present here a simple, readily accessible approach to identify monovalently tethered substrates for single-molecule enzymology and force spectroscopy, and apply our method to visualize single-molecule proteolysis in real time using a standard, inverted microscope equipped with magnetic tweezers (Figure 1). We capture beads onto chip-bound protease substrates in a microfluidic chamber, and computationally sort tracked substrate-tethered beads that are flow-stretched, showing that characteristic displacements under flow, or <u>f</u>low <u>ex</u>tension (FLEX) signatures, are a reliable method to identify beads tethered to a single substrate molecule. We then demonstrate specific proteolysis of bead-tethered substrates for two different classes of proteases. This technique should be a universal approach to sort putative substrates for a wide range of proteases, or other hydrolytic enzymes, and should be applicable to studies investigating proteolytic sensitivity as a function of force as well as to classic force-clamp spectroscopy experiments measuring the strength of adhesion bonds.

**Figure 1.**
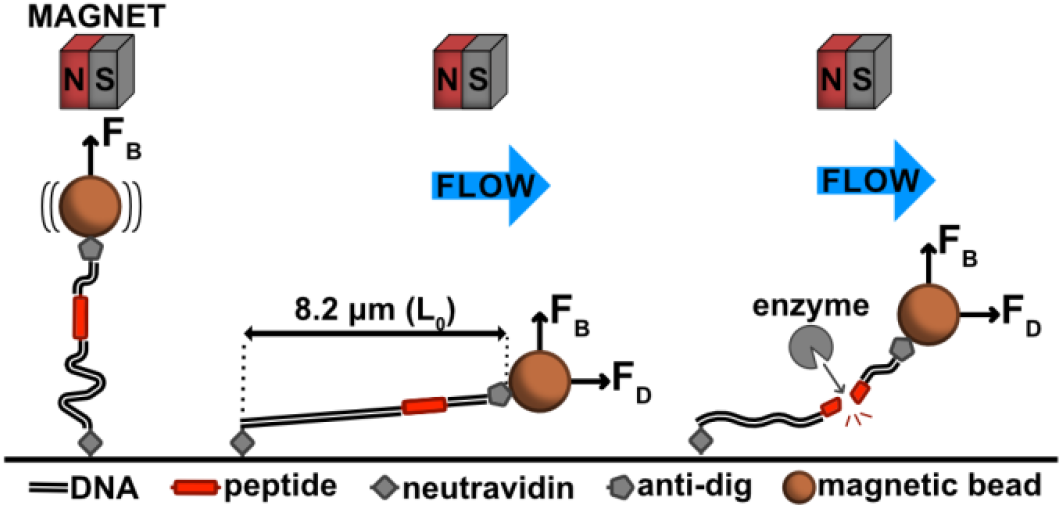
Schematic of the flow extension (FLEX) cleavage assay. Details of the FLEX approach are described in the text. F_B_, magnetic force; F_D_, drag force; L_0_, contour length.

## Results

In order to develop a reliable bead-tethered assay for single-molecule proteolysis, we developed an approach that relies on the use of flow extension (FLEX) to distinguish beads tethered to single substrates from beads with multiple tethers or beads that adhere to the flow-cell surface after capture. There are two independent measurements made in this experiment: first, we measure bead displacement in a flow extension assay to distinguish among different tether configurations, and then we measure bead release in a cleavage assay performed with protease (Figure 1). Our substrate contains a long DNA tether with an embedded polypeptide sequence between digoxigenin- and biotin-labeled DNA handles. The substrate is captured on a neutravidin surface using the biotinylated DNA end, and α-digoxigenin F_ab_ coated magnetic beads are then tethered to the beads at the digoxigenin-labeled end. This bead-tethered substrate is then used for multiplexed, single-molecule proteolysis (Figure 1).

The assembly of the DNA-conjugated peptide substrate is schematically illustrated in Figure 2. First, we link a single-stranded oligonucleotide to the C-terminal cysteine residue of a fluorescently labeled acceptor peptide using a maleimide-modified nucleotide (**1** in Figure 2A and Figure S1). We then couple our substrate peptide to this acceptor-DNA conjugate using evolved Sortase A (SrtA) (**2** in Figure 2A and Figure S1). Again using maleimide coupling, a second single-stranded oligonucleotide is attached to the N-terminal cysteine of this protein-DNA conjugate to generate a peptide substrate with single stranded oligonucleotides at each end (**3** in Figure 2A and Figure S1). After purification on a size exclusion column, this molecule is then ligated to a short DNA duplex at the N-terminal end, and to an XbaI-cleaved fragment of λ phage (23,998 bp) at the C-terminal end. Ligation of a digoxigenin-modified oligonucleotide to the short duplex, and a biotinylated oligonucleotide to the λ phage fragment results in production of a substrate with a biotin handle for surface attachment and a digoxigenin handle for capture of anti-digoxigenin beads (**4** in Figure 2B).

**Figure 2.**
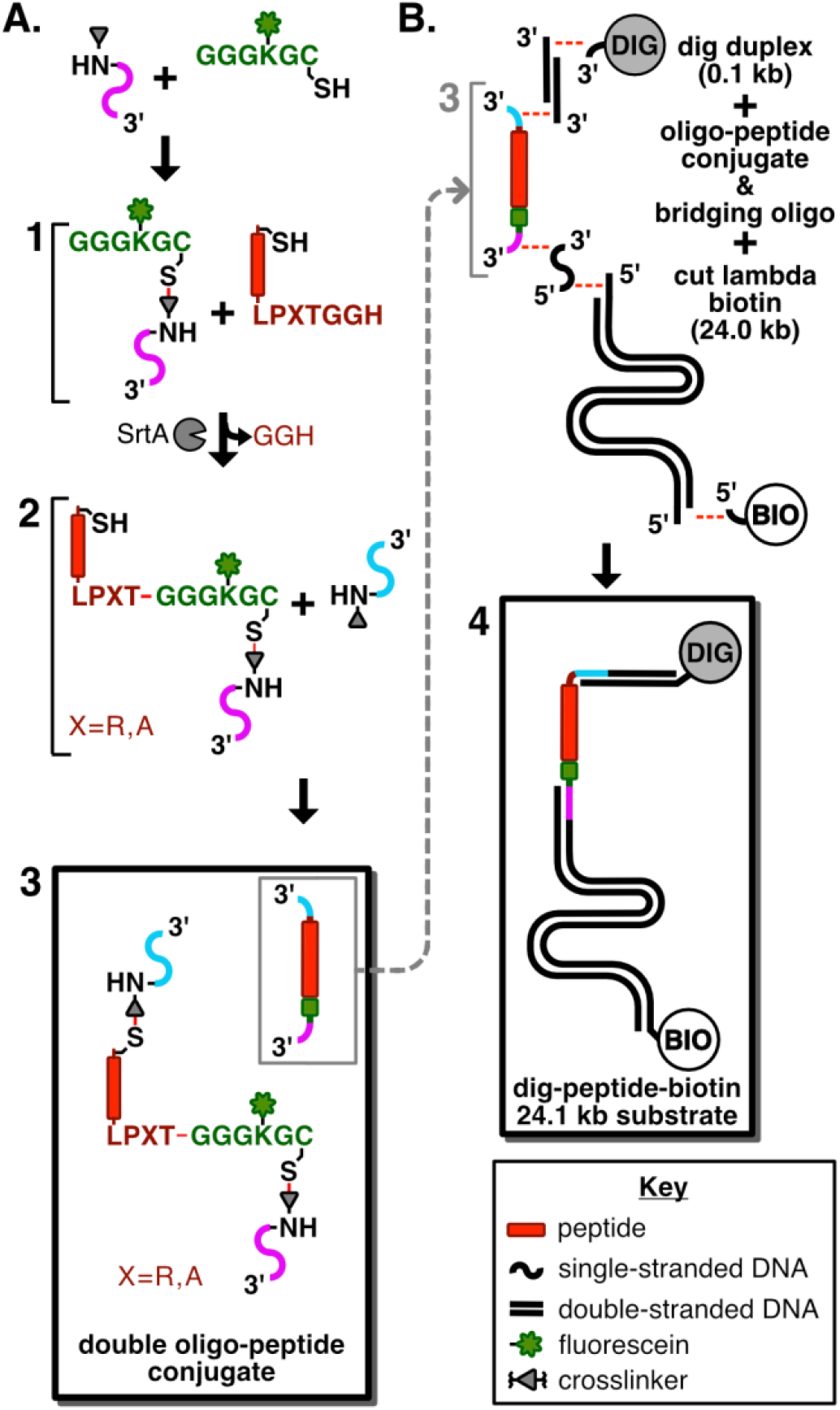
Scheme showing the route used to prepare DNA-conjugated peptide substrates for single-molecule proteolysis. A. Synthetic route to the double oligonucleotide-coupled peptide conjugate. Product **1** is a polyglycine conjugate made with a fluoresceinated lysine residue and a C-terminal thiol (green), coupled to a 5’-maleimide-oligo (magenta). Product **2** is the Sortase A-catalyzed conjugation of the substrate peptide containing a C-terminal LPXTG sortase acceptor motif (red) to the polyglycine-oligo (green-magenta). Product **3** (boxed) is the double oligonucleotide peptide conjugate made from linking the N-terminal cysteine of **2** to the 5’-maleimide-oligo (cyan). B. Synthesis of the digoxigenin-peptide-biotin substrate. Product **4** (boxed) is produced by an annealing and ligation reaction of a 5’ digoxigenin-containing oligonucleotide, a short (100 bp) DNA duplex and the N-terminal oligonucleotide end of the double oligo-peptide conjugate, along with simultaneous ligation of the C-terminal-end oligonucleotide to the XbaI-cleaved 24.0 kbp fragment of phage λ, and a terminal biotinylated oligonucleotide.

The final substrate for proteolysis contains a custom peptide embedded between DNA duplexes that together total 24,098 base-pairs and have a maximal extension length (L_0_) of 8.2 μm, assuming 0.34 nm per base pair and 0.35 nm per amino acid (Figures 1 and 2B). The modular nature of both N- and C-terminal DNA fragments enables the use of numerous surface attachment strategies and choice of DNA handles of variable lengths on either side of the substrate.

To establish proof-of-concept for the FLEX sorting method and optimize conditions for single-bead capture, we first investigated a control all-DNA tether of comparable length, assembled similarly (Figure S2). We investigated beads captured onto these control DNA tethers loaded onto the flow cell at four different concentrations, ranging from 0.2 – 25 pM (Figure 3A). At each concentration, we performed a FLEX experiment by measuring the position of individual beads in the absence and presence of flow to classify bead trajectories into immobile (extension <1.6 μm), partially mobile (extension between 1.6 – 4.8 μm), or fully mobile (extension between 4.8 – 8.7 μm), eliminating clumped beads or trajectories that were incomplete over the flow-extension period (Table S1). The trajectory data show that loading substrates at a concentration of 0.2 pM results in a majority of single beads with full trajectories falling into the fully mobile category, and that a concentration of 5 pM or greater results in multiple tethering or immobility of virtually all loaded beads (Figure 3C, D; see also Table S1). For the trajectories that are fully mobile, displacements fit to a normal distribution (Figure 3C and 3D, green population) of 6.37 ± 0.54 μm (mean ± S.D., n = 518). It is also clear that there is an inverse correlation between the fraction of fully mobile and immobile beads as the concentration of tethered beads increases, indicating that as more trajectories are observed there are sufficient bound substrates to cause multiple tethering for nearly every bead (Figure 3C and 3D). Overall, the bead mobility analysis indicates that the FLEX signature is a robust method to determine whether a bead is tethered to a single substrate molecule.

**Figure 3.**
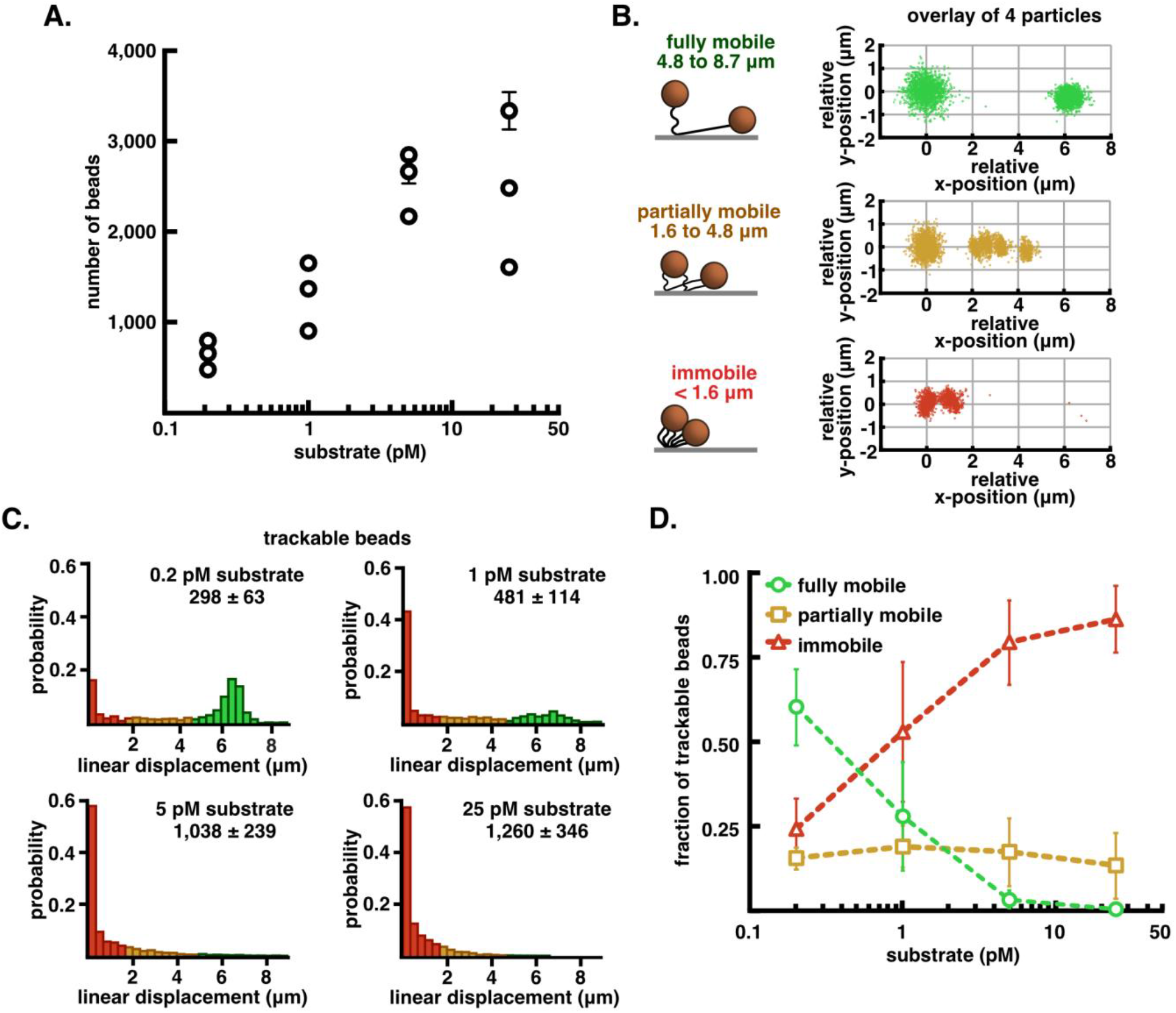
Tethered bead movement under flow extension as a function of tether density. A. Semi-log plot of the average number of captured beads in six fields-of-view from flow-chambers incubated with increasing amounts of dig-λ_XbaI_-biotin (0.2, 1, 5, 25 pM), reported as mean ± S.E.M. B. Graphic representation of the three classes of beads that are observed in every experiment alongside an overlay of relative initial (t = 0-200 sec) and extended (t = 300-350 sec) position data from four trajectories from each class of displacement: fully mobile (4.8 - 8.7 μm, green), partially mobile (1.6 – 4.8 μm, orange), and immobile (< 1.6 μm, red). C. Probability normalized histograms of the linear displacement of bead trajectories at increasing amounts of substrate. The number of particle trajectories tracked in 3 replicates (mean ± S.E.M.) are listed above each histogram. D. Quantification of the fraction of the total trajectories that were sorted into each class of displacement, reported as mean ± S.E.M. See also Table S1.

We next used our FLEX sorting procedure to assess the single-molecule proteolysis of a bead-tethered peptide substrate. For proof-of-concept studies, we assembled a double-DNA conjugated 19-residue peptide substrate (see Figure 2) containing a consensus Tobacco Etch Virus (TEV) protease cleavage site ^13^ (ENLYFQ/G, with the scissile bond indicated by a backslash), and subjected this substrate to cleavage with TEV (Figure 4). Beads captured on the DNA-peptide substrate show a distribution of FLEX behavior (Figure 4A,B), with 63% of the tethered beads (76/120) being fully mobile. Upon TEV treatment (Figure 4C), the population of tethered substrates (black) shows two kinetic phases and a residual population of approximately 45% of the beads that remain uncleaved after 10 min. Computational sorting of the beads into accepted and rejected beads based on their FLEX trajectories shows that the computationally rejected population (gray) resists specific cleavage by TEV, whereas the fully mobile population (green) shows more complete cleavage with a residual bead population of approximately 20% uncleaved beads after 10 min. The accepted bead population exhibits a rate of decay that is much faster than that of the rejected bead population (Figure 4C, compare gray and green decay curves). Whereas the accepted population of molecules for the specific peptide substrate shows rapid decay with a k_loss_ of 3.09 ± 0.03 × 10^−2^ sec^−1^ (n = 3, mean ± 95% CI), the accepted molecules for control bead-tethered substrates that lack peptide or tethered substrates treated with buffer under flow without added TEV exhibit only slow phases of bead loss that are nearly equivalent (k_loss_= 3.3 ± 0.2 × 10^−5^ sec^−1^ for the DNA-only control substrate, n = 3 mean ± 95% CI) likely attributable to dissociation of the anti-digoxigenin/digoxigenin (or neutravidin/biotin) interaction (Figure 4D).

**Figure 4.**
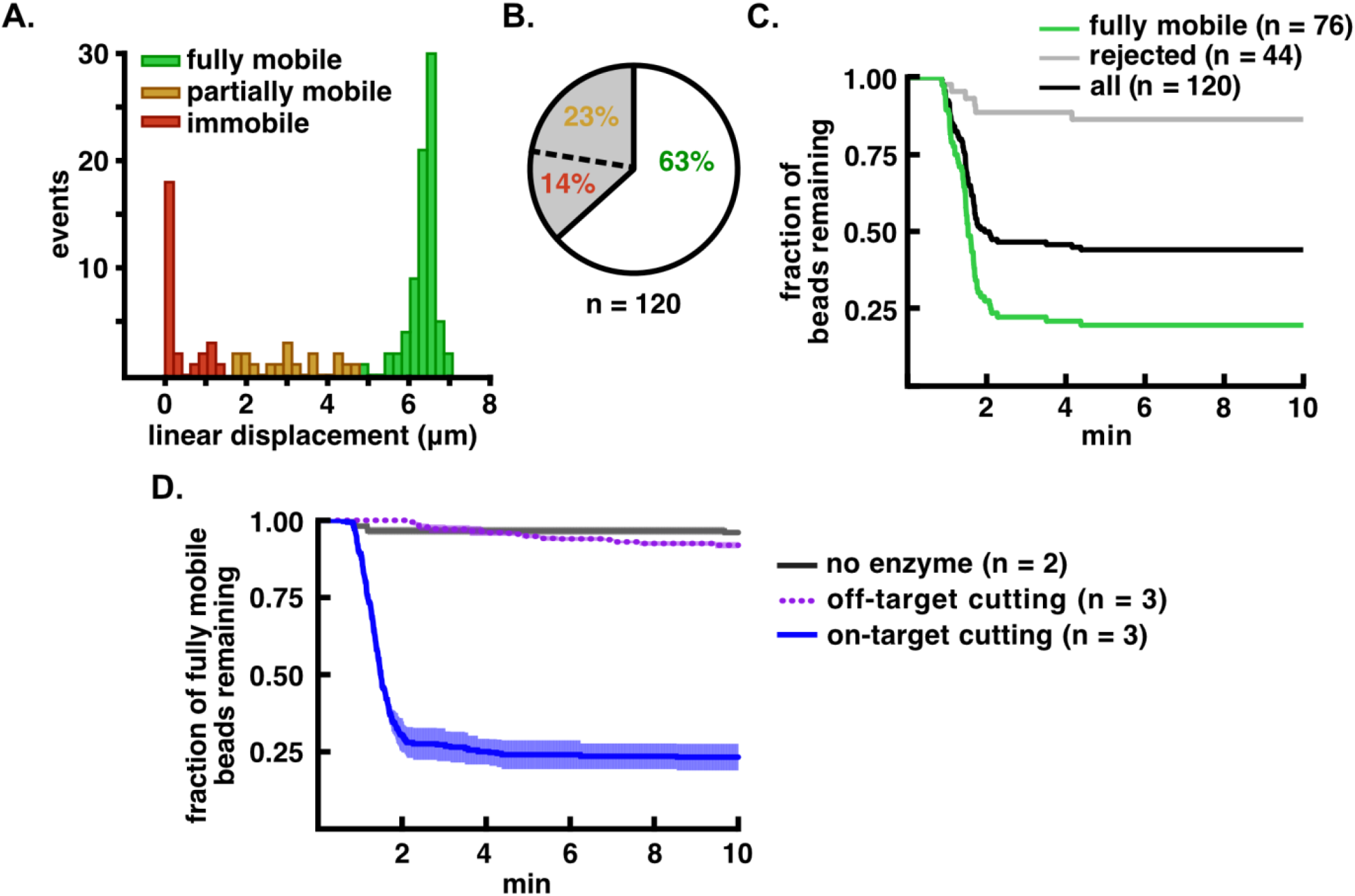
Improved analysis of TEV-catalyzed proteolysis using computational sorting of bead mobility. A. Distribution of flow extension distances for bead-tethered, TEV substrate molecules. The fully mobile population is shown in green, partially mobile in orange, and immobile in red. B. Pie chart showing the percentage of accepted, partially mobile, and immobile beads in the tethered population. C. Proteolysis of bead-tethered substrates upon addition of 20 μM TEV protease. The fraction of beads lost is plotted as a function of time for the total population (black), the accepted (fully mobile) population (green), and the rejected population (gray). D. Cleavage specificity analysis. Bead release from cleavage of the tethered peptide substrate is indicated with a blue solid line, release of the tethered all-DNA substrate is indicated with a purple dotted line, and bead release in the absence of enzyme is indicated in gray. Kinetic traces are normalized such that t = 0 min is set to 1 min before the start of bead loss, and are shown as mean ± S.E.M for the number of trials indicated (n = 2 or n = 3).

We next tested whether our assay system could be used to investigate proteolysis at the single-molecule level for a less selective protease, ADAM17. ADAM17 catalyzes the cleavage of a number of different proteins involved in intracellular signaling, including the epidermal growth factor precursor, tumor necrosis factor α, and, under certain pathophysiologic contexts, the force-sensitive substrate human Notch1 ^14–15^. Unlike TEV, the catalytic domain of ADAM17 does not recognize a highly specific consensus sequence, and is instead capable of cleaving a wide range of substrates with preference only for a bulky aliphatic residue in the P1’ position (i.e. Val or Ile) ^16–18^.

The ADAM17 substrate contains a ten-residue sequence from human Notch1 (NIPYKIEA/VQS, with the scissile bond indicated by a backslash) that spans the metalloprotease cleavage site (called S2 ^15, 19^) with additional short flanking sequences embedded between the N- and C-terminal DNA handles (Figure 2). Cleavage of the Notch1 substrate by ADAM17 shows both a fast phase of bead release dependent on the concentration of ADAM17 in the cleavage buffer, followed by a slow phase (Figure 5A). Controls testing for release of tethered beads in the absence of enzyme, for bead release in the presence of both ADAM17 and the metalloprotease inhibitor BB-94, or for release of beads tethered to a DNA-only substrate analog, show a slow phase of non-specific bead detachment, confirming that the fast exponential phase for ADAM17-catalyzed cleavage of the Notch1 substrate is due to specific single-molecule proteolysis. The rate of enzyme-catalyzed bead release plotted as a function of ADAM17 concentration shows saturable kinetics (Figure 5B), with an estimated k_max_ of 2.3 ± 0.2 × 10^−2^ and EC50 for [ADAM17] of 3.2 ± 0.7 × 10^−6^ M (reported as mean ± S.E.M., analogous to values for kcat and K_D_, as proposed previously ^10^).

**Figure 5.**
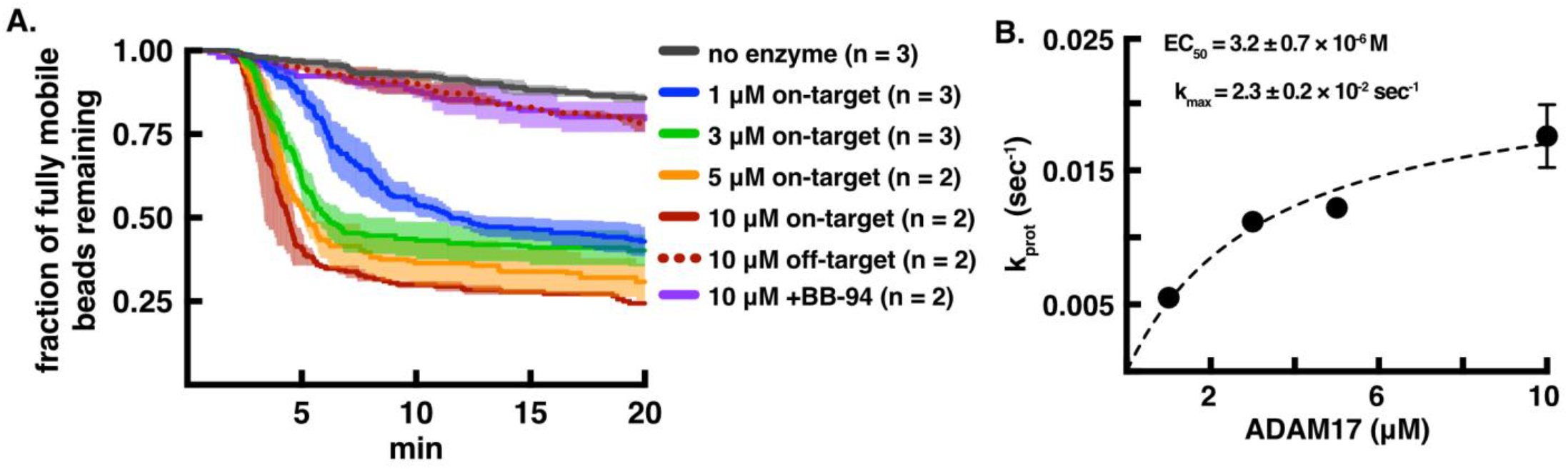
Single-molecule proteolysis of a Notch1 substrate by the ADAM17 catalytic domain. A. Plot of beads released as a function of time using different concentrations of enzyme and DNA control (dashed line) or Notch1 peptide substrate (solid lines). A cleavage reaction (at an ADAM17 concentration of 10 μM) performed in the presence of the metalloprotease inhibitor BB94 is shown in purple. B. Fit of the single-molecule cleavage rates from (A) as a function of ADAM17 concentration to the equation k_prot_ = k_max_ × [ADAM17]/(EC50 + [ADAM17]). Kinetic traces are shown as mean ± S.E.M for the number of trials indicated (n = 2 or n = 3).

## Discussion

Here, we report a method that uses flow extension sorting of tethered beads to enable stringent analysis of single-molecule proteolysis for two different enzymes and their substrates. The cleavage rates we observe are consistent with published studies of these enzymes in bulk proteolysis experiments ^20–21^. Our approach relies on computational sorting of substrate-tethered beads loaded at low density, using their mobility under flow to exclude immobile or multiply tethered beads and thereby only analyze beads tethered to a single substrate molecule.

In previous studies measuring proteolysis of bead-tethered substrates, approaches to classify different beads within the field of view were not used, and therefore it was not possible to make strong claims about monovalency ^6, 10^. Our data show that tether valency is highly sensitive to the concentration of substrate used for capture in the flow cell. Even when DNA-only molecules are used for the flow-extension analysis, a pronounced increase in fully mobile beads is readily evident as the decrease in grafting density is reduced to the lowest loading concentration (0.2 pM). The FLEX procedure enables empirical optimization of substrate concentration to maintain enough beads for multiplexing statistics while minimizing the population of beads rejected because of surface adsorption or multiple tethering. In the studies reported here, we were able to load all-DNA molecules into the chamber so that as many as 202 fully mobile (of 355 total) DNA-tethered beads were present in a single field of view. Even the double-oligonucleotide conjugate peptide substrates, which are likely to be more adsorption prone because of their embedded peptide sequences, can be delivered to the flow cell at concentrations sufficient to produce 120 fully mobile (of 165 total) substrate-tethered beads in a single field of view. We also note that our empirical data are consistent with statistical predictions about multiple tethering made using an ideal surface model dependent upon the grafting density of substrates and the sum of tether contour length and bead diameter ^22^.

The use of the flow-extension signature to perform computational exclusion of beads that are not held by a single tether results in a demonstrable improvement in the measurement of substrate cleavage kinetics (Figure 4). The approach should be particularly valuable in identifying reaction conditions suitable for investigation of protein or peptide substrates that are prone to adsorption on surfaces. Optimization of reaction conditions can be carried out by selecting blocking conditions and substrate variants that minimize the number of transiently or permanently adsorbed beads, enriching for the fully mobile beads that report faithfully on cleavage of a single substrate molecule during the enzyme-catalyzed reaction.

The assay tools reported here should be readily adaptable for the investigation of any tension dependent hydrolytic reaction, simply by using the magnetic tweezers to vary the magnitude of applied force. Apart from testing mechanosensing domains, our assay can be used to study the stability of peptide and foldamer secondary and tertiary structural elements by investigating the force-dependence of denaturation of such structural elements using proteolytic cleavage as a proxy. Two classes of molecules with changes in cleavage sensitivity can be envisioned: those substrates that have cryptic cleavage sites that are exposed upon denaturation or conformational change ^23–24^, or those substrates that only present a productive cleavage site when they fold^25–28^.

The FLEX sorting technique should also be applicable to *in vitro* single-molecule force-spectroscopy. For bond rupture experiments, it is equally important to ensure that only monovalent interactions are scored to evaluate catch- or flex-bond versus slip-bond behavior ^3, 29–30^.

## Materials and Methods

### Chemicals

Custom synthesized biopolymers were obtained from Genscript or IDT, and are listed in Table S2 in the Supporting Information. Sulfo-SMCC (sulfosuccinimidyl 4-(N-maleimidomethyl)cyclohexane-1-carboxylate; 22322), and APMA (4-aminophenylmercurial acetate; A9563) were from Sigma. BB-94 or batimastat was from Calbiochem (196440). NeutrAvidin (31000) was from Thermo Fisher. The anti-digoxigenein Fab was purchased from Roche (11214667001). Nano-strip was from KMG (210034). Key reagents for the single-molecule working buffer were bovine serum albumin, heat shock fraction (A7906) and Pluronic F-127 (P2443) from Sigma. M-280 Tosyl activated Dynabeads (14203) were from Invitrogen. SDS-PAGE analysis was performed with either Bolt 4-12% Bis-Tris Plus gels (Life Technologies, NW04125BOX) or 4-20% tris-glycine gels (Bio-Rad, 456-1096). Denaturing PAGE of oligonucleotides and their conjugates was performed with Novex 15% TBE-Urea gels (Life Technologies, EC6885BOX). PD-10 desalting columns were from GE (95017-001). P-6 (7326221) and P-30 (7326223) Micro Bio-Spin columns were obtained from Bio-Rad.

Lambda (λ) DNA dam(-) was purchased from New England Biolabs (N3013L). 3-aminopropyltrimethoxysilane was purchased from Sigma (281778-100ML). Methyl-PEG5000-succinimidyl valerate (mPEG5000-SVA) and biotin-PEG5000-SVA (bioPEG5000SVA) were acquired from Laysan Biosciences and stored dessicated at −20 C.

### Enzymes

T4 polynucleotide kinase (M0201S), T4 DNA ligase (M0202S), XbaI (R0145L), and XhoI (R0146S) were obtained from New England Biolabs. The enhanced activity SrtA pentamutant was expressed and purified from bacteria using the pET29-eSrtA vector (Addgene, #75144) as previously reported ^31^. TEV protease was also expressed recombinantly in bacteria using the pTrc-7H-PRO plasmid and purified as described ^32^. The catalytic domain of ADAM17 (residues 215-477) was isolated after recombinant expression of its precursor form (1-477) with a C-terminal hexahistidine tag using a baculovirus expression system in insect cells. Purification was carried out in the presence of APMA as described ^33^. Enzyme activity was confirmed in bulk solution using a fluorogenic substrate (Mca-PLAQAV-Dpa-RSSSR-NH_2_; R&D systems, ES003) ^21^. Enzymes were snap frozen and stored at −80 C in aliquots before use.

### Custom synthesized biopolymers

Oligonucleotides listed in Supplementary Table S2 were custom synthesized from Integrated DNA Technologies (IDT). Peptides listed in Supplementary Table S2 were custom synthesized from Genscript and supplied at 95% minimum purity. SMCC-activated amine-oligonucleotides (activated oligonucleotides, Table S2) were produced by mixing sulfo-SMCC (7.5 mM, dissolved in dimethylformamide with 300 μM amino-oligonucleotide at 25° C in 10 mM phosphate buffer, pH 7.2, containing 150 mM NaCl (PBS). After a 3-5 h reaction time, the activated oligonucleotide was purified away from residual free crosslinker on a PD-10 desalting column(s) pre-equilibrated in PBS, concentrated on a 3kDa MWCO centrifugal filter, and used immediately for peptide conjugation.

### Other materials

Tubing was PE60 (Stoelting, 51162), coverslips were No. 1.5 VistaVision (VWR, 16004-312), double sided tape (Grace Bio-Labs, 620001), and 1 mm quartz tops were from Technical Glass Products.

### Preparation of coverslips and flow chambers

No. 1.5 coverslips were prepared by first etching with stabilized piranha solution for 2 h, then silanized with pre-hydrolyzed 3-(aminopropyl)trimethoxysilane in acidic methanol, and passivating by alkylation with PEG_5000_-succinimide containing 2-6 mol% biotinylated PEG ^34–35^. Coverslips were stored under vacuum before use. To generate flow cells, 15 × 5 mm channels were cut into double-sided tape, the tape was placed between the coverslip and a glass slide, and inlet and outlet ports were attached to the glass with epoxy as described ^34, 36^.

### Synthesis of double oligonucleotide peptide conjugates

The oligo-peptide conjugate (**1**) (Figure 2; oligo-peptide conjugate 1, Table S2) was synthesized by conjugation of the H_2_N-GGGK*GC-COOH (where the K* residue is ε-[5-amino-fluorescein (5-FAM)]-Lys) sortase donor peptide to a 5’-maleimide activated 17-base pair oligonucleotide (activated oligonucleotide 1 and peptide-oligo conjugate 1, Table S2) ^37^. The conjugation reaction was allowed to proceed for 3 h at 22-25°C in PBS buffer pH 7.2 (10 mM phosphate, 150 mM NaCl) with 1 mM TCEP, containing 2 mM (~15 eq.) peptide 1 (Table S2) and 130 μM (1 eq) activated oligonucleotide 1 (Table S2). To purify the conjugate (**1**) (oligo-peptide conjugate 1, Table S2) from free peptide, the reaction was exchanged into in HBS buffer (25 mM HEPES pH 7.5, 150 mM NaCl) with a gravity desalting column and purified by size-exclusion chromatography to apparent homogeneity as assessed by TBE-Urea PAGE. The conjugate was stored at −80°C until use.

To produce the oligo-peptide conjugate (**2**) (Figure 2; oligo-peptide conjugates 2-N and 2-T, Table S2), conjugate (**1**) was coupled to the C-terminus of either the Notch S2 peptide (peptide 2, Table S2) or the TEV substrate peptide (peptide 3, Table S2) in a transpeptidation reaction using SrtA. The SrtA transpeptidation was performed over 4 h at 42°C in HBS pH 7.5 with 10 mM MgCl_2_ and 4 μM NiCl_2_ reaction buffer using 15 μM (1 eq.) of oligo-peptide conjugate 1 (**1**), 150 μM (10 eq.) peptide 2 or 3, and 15 μM (1 eq.) eSrtA, similar to previously reported methods ^38–39^. After coupling, the conjugate (**2**) was further purified using a desalting column, which also was used for buffer exchange into HBS [25 mM HEPES pH 7.5, 150 mM NaCl]. Purity was assessed by TBE-Urea PAGE analysis. Conjugate (**2**) was stored at −80°C until use.

The double oligo-peptide conjugate (**3**) was formed by attaching the N-terminal cysteine of conjugate (**2**) to a 5’-maleimide functionalized 20 base-pair oligonucleotide (activated oligonucleotide 2, Table S2), essentially as described above for synthesis of conjugate (**1**) except that activated oligonucleotide 2 was used in vast molar excess during this step. The Notch S2 and TEV double oligo-peptide conjugates (**3**) (Figure 2; oligo-peptide conjugates 3-N and 3-T, respectively, Table S2) were purified using size-exclusion chromatography to apparent homogeneity, as assessed by denaturing TBE-Urea PAGE, and were stored at −80°C.

### Production of duplex DNA single-molecule substrates

Each double-oligonucleotide peptide conjugate (**3**) was annealed to a bridging oligonucleotide (oligonucleotide 3, Table S2), a 100 base-pair 5’-digoxigenin containing sequence from the pUC19 *ori* region (oligonucleotides 4, 5, and 6; Table S2), an XbaI digested fragment from the phage λ genome (λ_XbaI_, 23,998 base-pairs, 3.75 nM, at 250 μg scale), and a 3’-biotin cohesive site cosR sequence (oligonucleotide 7, Table S2) in 20 mM HEPES buffer, pH 7.5 (annealing temperatures and times were similar to previously published methods ^40^). Prior to annealing, all annealed oligonucleotides with free 5’ ends were treated with T4 polynucleotide kinase in 1X PNK buffer for 1 hr at 37°C and purified with a Qiagen nucleotide cleanup kit). The annealed assembly was ligated with 0.16 U/μL T4 DNA ligase for 2 hr at room temperature. The fully ligated substrate was purified away from lower molecular weight contaminants (removal of the free biotinylated oligonucleotide is critical) by buffer exchange into HEPES-buffered saline (HBS), pH 7.5, using a centrifugal filter with a molecular weight cutoff of 100 kDa. The all-DNA control substrate was produced using the same assembly and ligation procedure, using a DNA insert (oligonucleotide 8, Table S2) in place of the double oligonucleotide conjugate.

### Substrate capture and bead tethering procedure

Flow chambers were mounted on the stage of an Olympus IX51 inverted microscope on an air table (Technical Manufacturing Corporation), with off-axis LED gooseneck illuminator (Fisher), and custom-built magnetic tweezers, as previously described with minor modifications ^6^. The chambers were also equipped with a syringe pump (Harvard Apparatus) controlled using a stopcock valve and a 50 mL air spring to regulate fluid flow through the flow-cell. All images were acquired with a QIClick-R-F-M-12 Mono CCD camera (QImaging) controlled with Micro-Manager software ^41^.

Prior to capture of substrates on the flow-cell surface, the flow-cell chamber was first treated for a minimum of 10 min with 500 μL blocking buffer (HBS containing 0.5% (w/v) BSA and 1% (w/v) Pluronic F127) drawn manually into the flow cell. Then, the chamber was incubated for 3-5 min with 80 μL of NeutrAvidin (0.1 mg/mL) drawn manually into the chamber, and washed with 500 μL working buffer (HBS containing 0.1% (w/v) BSA and 1% (w/v) Pluronic F127) at a flow-rate of 100 μL/min. Substrate was diluted into working buffer (0.15 to 15 pM final concentration), and a total volume of 100 μL of substrate was drawn into the chamber first at 20 μL/min for one min to clear the inlet tubing, and at 5 μL/min thereafter before incubating in the chamber for an additional 2-5 min. The chamber was then washed with working buffer for 10 min at a flow rate of 50 μL/min.

To prepare beads for capture, tosyl-activated 2.8 μm Dynabeads were conjugated to anti-digoxigenin F_ab_, and stored in PBS with 0.01% (w/v) BSA at 4°C, as described (according to manufacturer’s Protocol and Tanner, N. and van Oijen, A. (2010) ^34^). Immediately before use, beads were diluted to 10 μg/mL in working buffer, vortexed, and sonicated to minimize clumping. The suspended beads (~250 μL) were then drawn into the chamber at 20 μL/min. The chamber was aligned in the direction of flow to make sure that a bead in motion across the chamber did not move more than half its diameter across the entire field of view (fov). The chamber was then washed with 400 μL of working buffer at 30 μL/min. After the washing step, the flow was halted to allow beads to relax from their extended position (typically for 5-10 mins). After the syringe pump was turned off, a 5mm NdFeB cube magnet on a hinged mount was positioned at 9 mm above the coverglass with a micrometer (equivalent to approximately 0.6 pN of applied force normal to the coverslip) in order to keep the beads above the surface. Force was estimated with a digoxigenin-λ-biotin (48,502 bp, full length prepared as described in ^40^) tether based on the variance in bead position (〈*σ_y_*^2^〉) in the direction transverse to applied force, according to the method published by Strick and colleagues ^42^.

### Single-molecule flow-extension procedure

For proof-of-concept flow extension experiments (Figure 3, supplementary movie V1), additional buffer was added to the inlet after substrate loading and bead capture, and the valve was opened and closed to allow pressure equilibration of the flow cell. Data acquisition was then initiated at 2 frames per second, initially in the absence of flow, but with the magnet in position (Figure 1), to allow determination of the initial position of each tracked bead. At the time of valve opening (at t = 200 sec), the buffer was drawn into the chamber at 5 μL/min using a syringe pump to induce flow-extension. The extension under flow reaches equilibrium at approximately 4 min. At the 5 min time point, the position of the bead at extension was recorded.

### Analysis and sorting of flow extension bead profiles

Time-lapse data sets were analyzed with custom MATLAB scripts written for this work, available upon request. Detected particles were identified and analyzed with colloidal particle tracking algorithms developed by Cocker, J.C. and Grier, D.G. ^43^ that were adapted for MATLAB by Kilfoil, M. ^44^. The trajectories of monodisperse particles were inspected manually to ensure high-quality tracking output, using the msdanalyzer suite created by Tinevez, J.-Y. ^45^.

The flow-extension trajectory of each monodisperse bead was analyzed to determine its linear displacement. Specifically, only bead trajectories that started at the first frame (t = 0 sec) of acquired data and that persisted through the time required for bead extension (t = 200 sec) were scored. The mean initial (0 to 200 sec) and extended (300 to 350 sec) positions were determined by fitting the bead positions to a Gaussian distribution. Linear flow-displacements were calculated by measuring the distance between these mean positions. The displacements were measured to sub-pixel accuracy (*σ_σ_RMSE__*) and precision 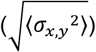 ^22^.

Based on their mean displacements, the analyzed beads were sorted into accepted (fully mobile [8.7 to 4.8 μm]) and rejected (partially mobile [4.8 to 1.6 μm] and immobile [1.6 to 0 μm]) populations. The histogram of displacements for a population of tethered substrates was always inspected to qualitatively assess the character of the distribution relative to the fraction in each population. The mean extension for a high-quality fully mobile population was typically between 6.0 – 6.5 μm (L/L_0_ = 0.73 – 0.79) assuming L_0_ = 8.20.

### Single-molecule proteolysis with TEV

Immediately before use, TEV protease was buffer-exchanged into TEV cutting buffer (25 mM HEPES pH 7.5, 25 mM NaCl, and 0.01%(w/v) Pluronic F-127) with a P-6 spin column to remove residual TCEP present during storage, and then diluted to a concentration of 20 μM before introduction into the flow cell. After substrate loading and bead capture, TEV was introduced into the flow cell at 5 μL/min and bead trajectories were monitored for assignment into mobile, partially mobile, or immobile categories. The field of view was then monitored for bead loss as a function of time to produce the cleavage plots in Figure 4 (see also supplementary movie V2).

### Single-molecule proteolysis with ADAM17

ADAM17 was diluted to the appropriate concentration into metalloprotease cutting buffer (HBS with 4 μM ZnCl_2_ and 1%(w/v) Pluronic F-127) immediately before use, and flowed into the chamber to generate the cleavage plots in Figure 5. Flow of enzyme into the chamber was halted after ~25 min, but trajectories were tracked for a total of 45 min to extract the rate constant for the slow exponential phase. To confirm that cleavage was due to ADAM17, the metalloprotease inhibitor BB-94 was used at 20 μM.

### Curve-fitting for cleavage data

Bead loss curves were fit to a double exponential using non-linear least squares fitting, where the fast rate constant corresponds to k_prot_. The start of bead loss was determined for each curve by manual inspection. For bead loss curves measured in the absence of enzyme or with inhibitor present, fits were to a single exponential over a time course of 15 min. For ADAM17 experiments, a plot of k_prot_ as a function of enzyme concentration was fit to:

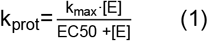

to extract k_max_ and EC50, which are analogous to k_cat_ and K_D_, as described previously ^10^.

## ASSOCIATED CONTENT

The Supporting Information is available free of charge on the ACS Publications website, and includes the following:

1. Supporting Figures S1 and S2, Supporting Tables S1 and S2, Legends for Supporting Movies V1 and V2, combined into a single PDF file.
2. Supporting Movies V1 and V2 (file type AVI)

## AUTHOR INFORMATION

### Corresponding Author

*To whom correspondence should be addressed.

### Author Contributions

Project conception: all authors; experimental design: all authors; data acquisition, A.A.D.; data interpretation and analysis: all authors; manuscript preparation: all authors; funding: S.C.B. and J.J.L.

## ACKNOWLEDGMENTS

The authors are grateful to Tom C. M. Seegar for providing the baculovirus used for ADAM17 production, Andrew Moreno for help with coding, and both Thomas G. W. Graham and Dan Song for help with flow cell preparation. This work is supported by NIH R35 CA220340 (to S.C.B.) and R01 GM114065 (to J.J.L.).

## Supporting Information

**Table S1.**
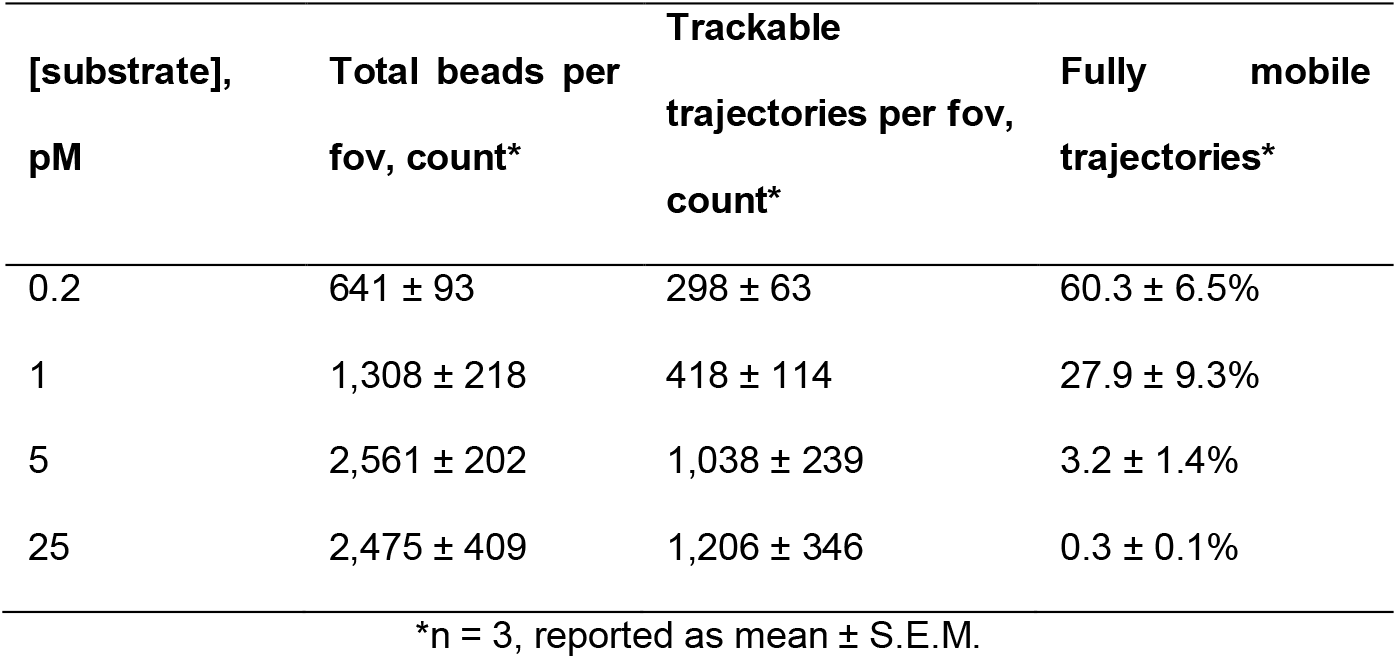
Relationship between DNA substrate binding density and trajectory mobility.

**Table S2.**
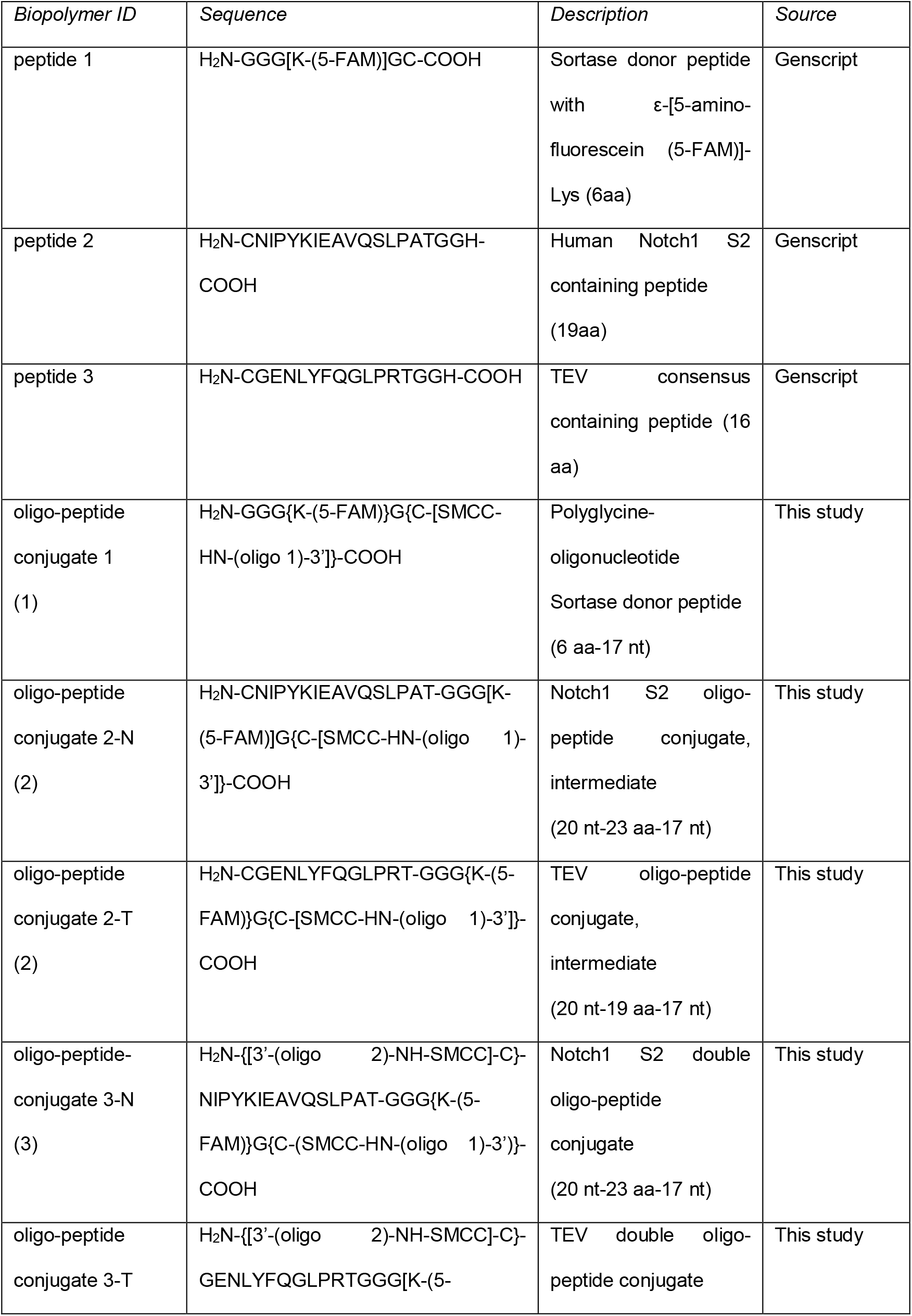
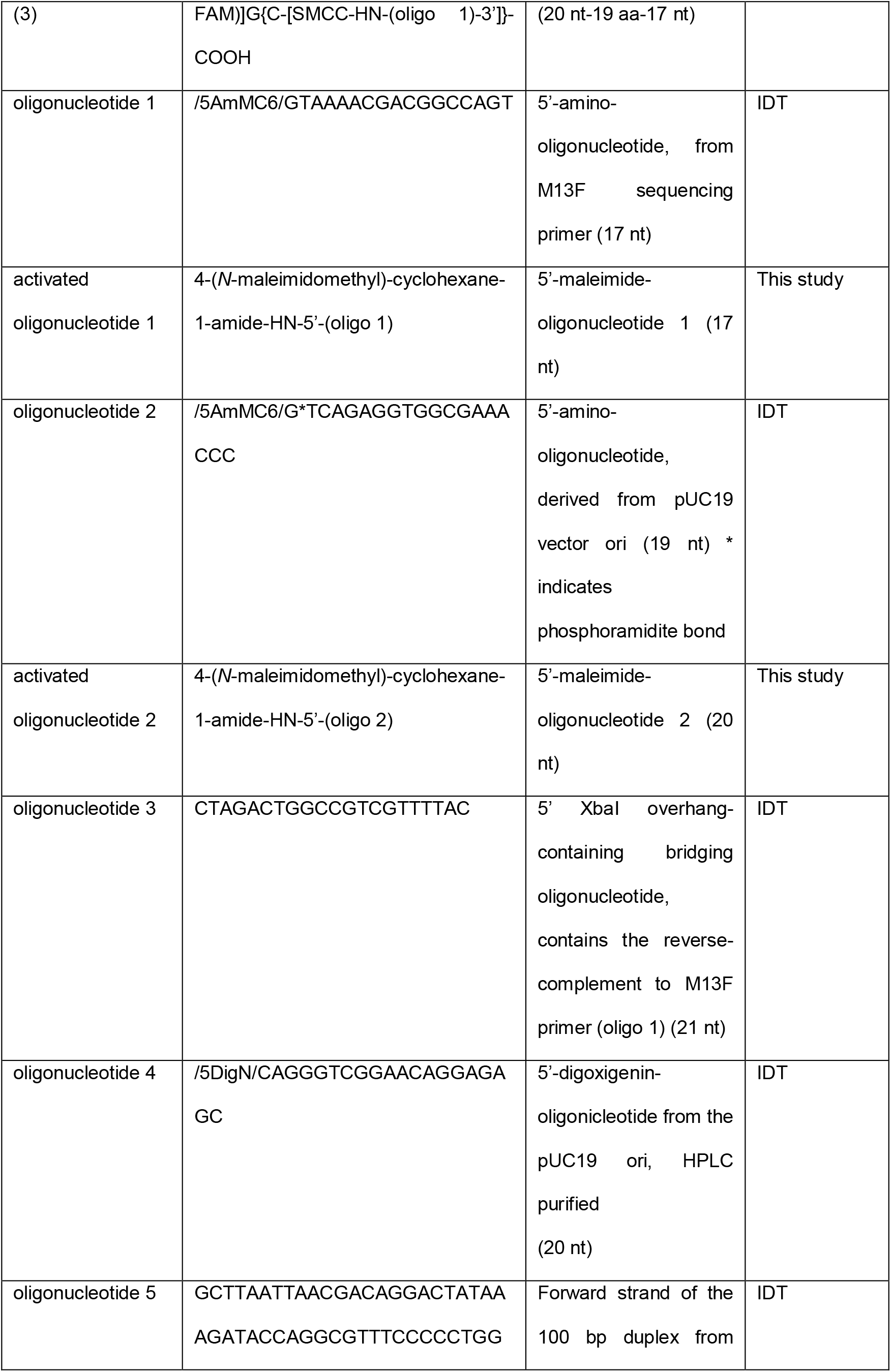
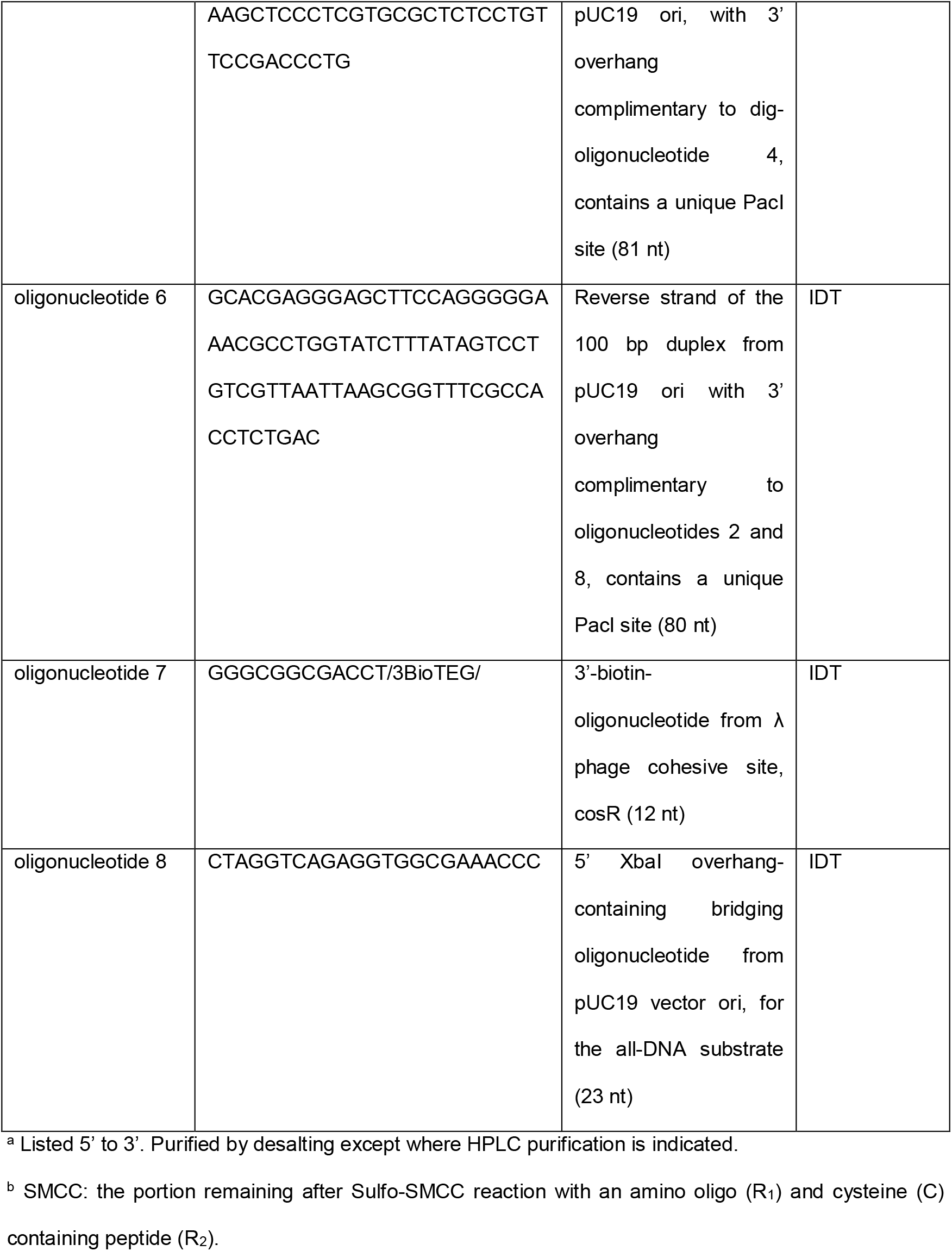
Peptides, oligonucleotides^a^, oligonucleotide-peptide conjugates^b^, and fluorogenic peptides used in this study.

**Figure S1.**
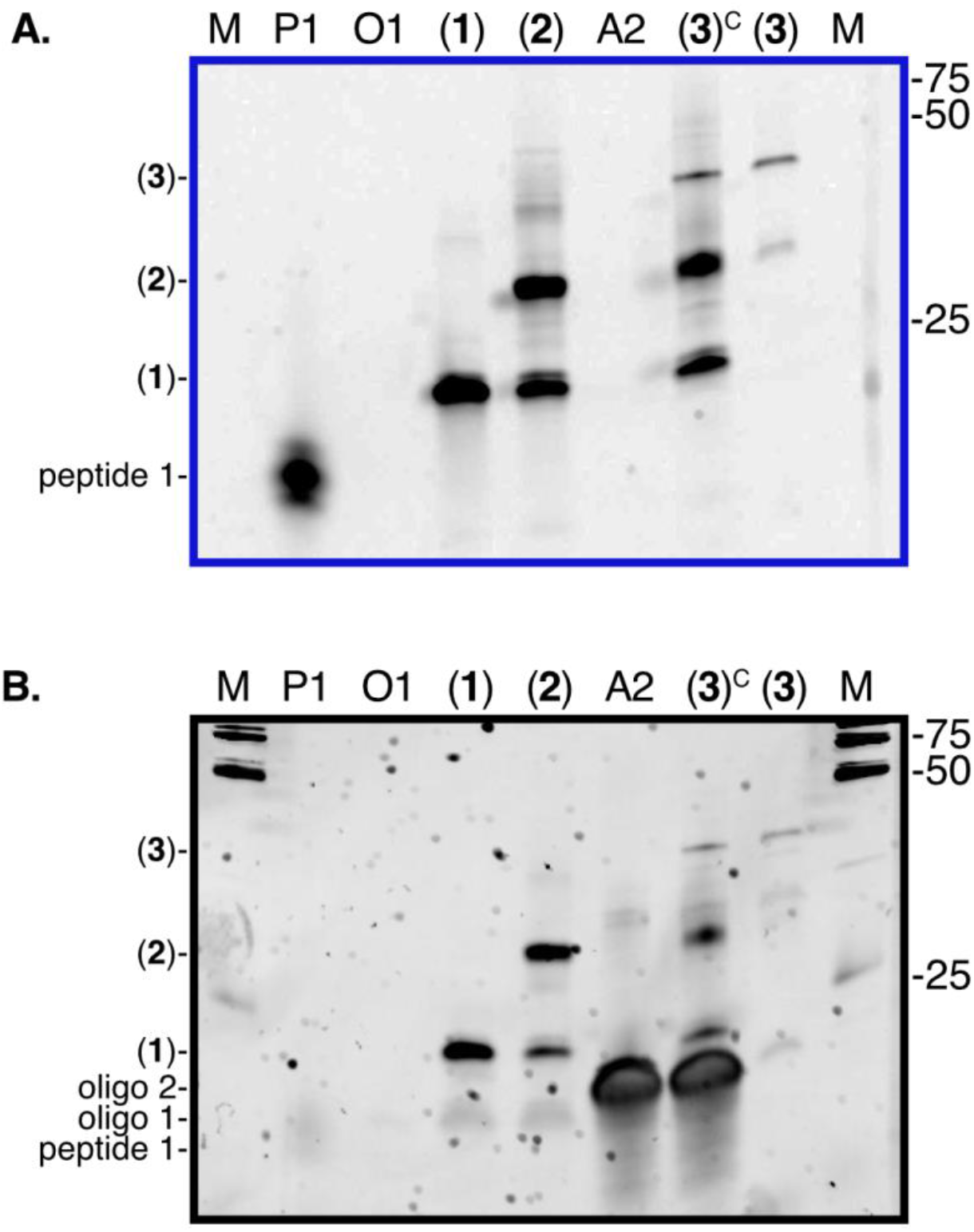
SDS-PAGE analysis of the double oligo-peptide conjugate intermediate for TEV cleavage. 15% TBE-Urea PAGE analysis showing: (A) in gel fluorescence of an unstained gel to detect the fluorescein label, and (B) in gel UV excitation fluorescence, of a SybrSafe stained gel to enable DNA detection. Abreviations above lanes: M, low MW DNA marker (NEB); P1, peptide 1; O1, oligonucleotide 1; (**1**) oligo-peptide conjguate 1; (**2**) partially purified oligo-peptide conjugate 2-T; A2, activated oligonucleotide 2; (**3**)^C^ crude double oligo-peptide conjugate 3-T; (**3**) double oligo-peptide conjugate 3-T.

**Figure S2.**
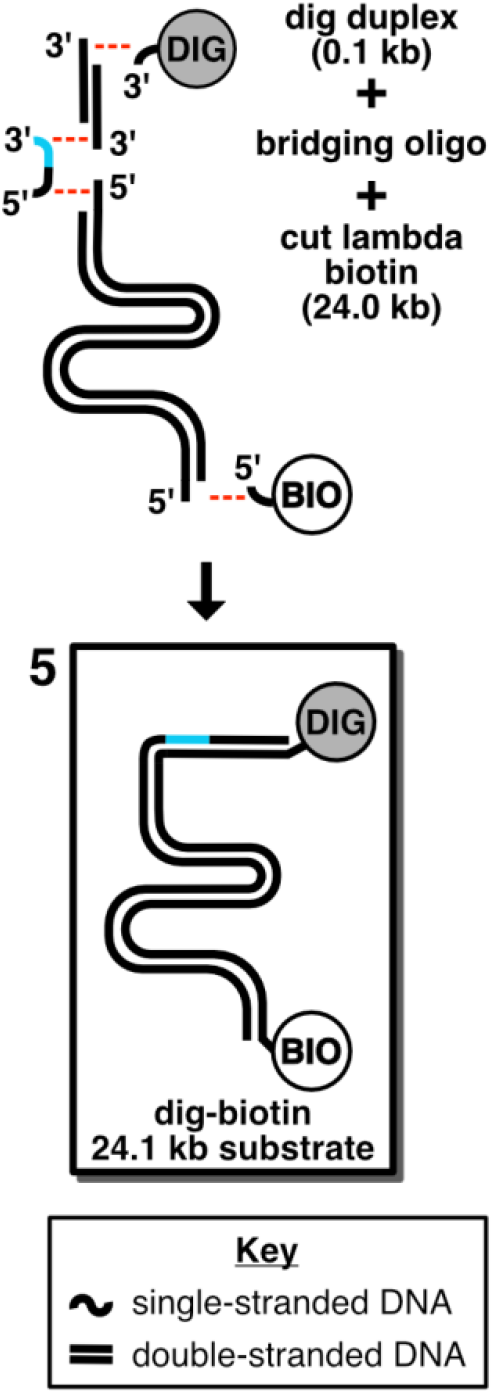
Scheme for assembly of DNA-only control substrate. Product (5) (boxed) is produced by annealing and ligation of a 5’ digoxigenin-containing oligonucleotide, a short (100 bp) DNA duplex, a bridging oligonucleotide (oligonucleotide 8, cyan and black), the XbaI-cleaved 24.5 kbp fragment of phage λ, and a terminal biotinylated oligonucleotide.

### Legends for Supporting Movies V1 and V2

**Supporting Movie V1. Flow extension of all-DNA substrate (5) captured at 0.2 pM**. The frame rate is 45 fps. Scale bar is 10 microns. Flow is from left to right. Flow starts at ~8 sec.

**Supporting Movie V2. Single molecule proteolysis of TEV consensus DNA-peptide conjugate (4) with TEV protease**. The frame rate is 45 fps. Scale bar is 10 microns. Flow is from left to right. Flow starts at ~8 sec. Cleavage starts at ~16 sec.

## References

1. Ingber, D. E., Tensegrity: the architectural basis of cellular mechanotransduction. Annu Rev Physiol 1997, 59 (1), 575–599.

2. Bustamante, C.; Bryant, Z.; Smith, S. B., Ten years of tension: single-molecule DNA mechanics. Nature 2003, 421(6921), 423–7.

3. Liu, B.; Chen, W.; Zhu, C., Molecular force spectroscopy on cells. Annu Rev Phys Chem 2015, 66, 427–51.

4. Sachs, F., Mechanical Transduction and the Dark Energy of Biology. Biophys J 2018, 114 (1), 3–9.

5. Zhang, X.; Halvorsen, K.; Zhang, C. Z.; Wong, W. P.; Springer, T. A., Mechanoenzymatic cleavage of the ultralarge vascular protein von Willebrand factor. Science 2009, 324 (5932), 1330–4.

6. Gordon, W. R.; Zimmerman, B.; He, L.; Miles, L. J.; Huang, J.; Tiyanont, K.; McArthur, D. G.; Aster, J. C.; Perrimon, N.; Loparo, J. J.; Blacklow, S. C., Mechanical Allostery: Evidence for a Force Requirement in the Proteolytic Activation of Notch. Dev Cell 2015, 33 (6), 729–36.

7. Langridge, P. D.; Struhl, G., Epsin-Dependent Ligand Endocytosis Activates Notch by Force. Cell 2017, 171 (6), 1383–1396 e12.

8. Saxena, M.; Changede, R.; Hone, J.; Wolfenson, H.; Sheetz, M. P., Force-Induced Calpain Cleavage of Talin Is Critical for Growth, Adhesion Development, and Rigidity Sensing. Nano Lett 2017, 17 (12), 7242–7251.

9. Vilfan, I. D.; Lipfert, J.; Koster, D. A.; Lemay, S. G.; Dekker, N. H., Magnetic Tweezers for Single-Molecule Experiments. In Handbook of Single-Molecule Biophysics, 2009; pp 371–395.

10. Adhikari, A. S.; Chai, J.; Dunn, A. R., Mechanical load induces a 100-fold increase in the rate of collagen proteolysis by MMP-1. J Am Chem Soc 2011, 133 (6), 1686–9.

11. De Vlaminck, I.; Henighan, T.; van Loenhout, M. T.; Pfeiffer, I.; Huijts, J.; Kerssemakers, J. W.; Katan, A. J.; van Langen-Suurling, A.; van der Drift, E.; Wyman, C.; Dekker, C., Highly parallel magnetic tweezers by targeted DNA tethering. Nano Lett 2011, 11 (12), 5489–93.

12. Yang, D.; Ward, A.; Halvorsen, K.; Wong, W. P., Multiplexed single-molecule force spectroscopy using a centrifuge. Nat Commun 2016, 7, 11026.

13. Carrington, J. C.; Dougherty, W. G., A viral cleavage site cassette: identification of amino acid sequences required for tobacco etch virus polyprotein processing. Proc Natl Acad Sci U S A 1988, 85 (10), 3391–3395.

14. Moss, M. L.; Jin, S. L.; Milla, M. E.; Bickett, D. M.; Burkhart, W.; Carter, H. L.; Chen, W. J.; Clay, W. C.; Didsbury, J. R.; Hassler, D.; Hoffman, C. R.; Kost, T. A.; Lambert, M. H.; Leesnitzer, M. A.; McCauley, P.; McGeehan, G.; Mitchell, J.; Moyer, M.; Pahel, G.; Rocque, W.; Overton, L. K.; Schoenen, F.; Seaton, T.; Su, J. L.; Becherer, J. D.; et al., Cloning of a disintegrin metalloproteinase that processes precursor tumour-necrosis factor-alpha. Nature 1997, 385(6618), 733–6.

15. Brou, C.; Logeat, F.; Gupta, N.; Bessia, C.; LeBail, O.; Doedens, J. R.; Cumano, A.; Roux, P.; Black, R. A.; Israël, A., A novel proteolytic cleavage involved in Notch signaling: the role of the disintegrin-metalloprotease TACE. Molecular Cell 2000, 5 (2), 207–216.

16. Caescu, C. I.; Jeschke, G. R.; Turk, B. E., Active-site determinants of substrate recognition by the metalloproteinases TACE and ADAM10. Biochem J 2009, 424 (1), 79–88.

17. Mohan, M. J.; Seaton, T.; Mitchell, J.; Howe, A.; Blackburn, K.; Burkhart, W.; Moyer, M.; Patel, I.; Waitt, G. M.; Becherer, J. D., The Tumor Necrosis Factor-α Converting Enzyme (TACE): A Unique Metalloproteinase with Highly Defined Substrate Selectivity. Biochemistry 2002, 41 (30), 9462–9469.

18. Tucher, J.; Linke, D.; Koudelka, T.; Cassidy, L.; Tredup, C.; Wichert, R.; Pietrzik, C.; Becker-Pauly, C.; Tholey, A., LC-MS based cleavage site profiling of the proteases ADAM10 and ADAM17 using proteome-derived peptide libraries. J Proteome Res 2014, 13 (4), 2205–14.

19. Mumm, J. S.; Schroeter, E. H.; Saxena, M. T.; Griesemer, A.; Tian, X.; Pan, D.; Ray, W. J.; Kopan, R., A ligand-induced extracellular cleavage regulates γ-secretase-like proteolytic activation of Notch1. Molecular Cell 2000, 5 (2), 197–206.

20. Stawikowska, R.; Cudic, M.; Giulianotti, M.; Houghten, R. A.; Fields, G. B.; Minond, D., Activity of ADAM17 (a disintegrin and metalloprotease 17) is regulated by its noncatalytic domains and secondary structure of its substrates. J Biol Chem 2013, 288 (31), 22871–9.

21. Minond, D.; Cudic, M.; Bionda, N.; Giulianotti, M.; Maida, L.; Houghten, R. A.; Fields, G. B., Discovery of novel inhibitors of a disintegrin and metalloprotease 17 (ADAM17) using glycosylated and non-glycosylated substrates. J Biol Chem 2012, 287 (43), 36473–87.

22. Yin, H.; Landick, R.; Gelles, J., Tethered particle motion method for studying transcript elongation by a single RNA polymerase molecule. Biophysical journal 1994, 67 (6), 2468–2478.

23. Stanger, H. E.; Gellman, S. H., Rules for Antiparallel β-Sheet Design: D-Pro-Gly Is Superior to L-Asn-Gly for β-Hairpin Nucleation1. J Am Chem Soc 1998, 120 (17), 4236–4237.

24. Cline, L. L.; Waters, M. L., The structure of well-folded beta-hairpin peptides promotes resistance to peptidase degradation. Biopolymers 2009, 92 (6), 502–7.

25. Zasloff, M., Magainins, a class of antimicrobial peptides from Xenopus skin: isolation, characterization of two active forms, and partial cDNA sequence of a precursor. Proc Natl Acad Sci U S A 1987, 84 (15), 5449–5453.

26. Resnick, N. M.; Maloy, W. L.; Guy, H. R.; Zasloff, M., A novel endopeptidase from Xenopus that recognizes α-helical secondary structure. Cell 1991, 66 (3), 541–554.

27. Bayer, P.; Arndt, A.; Metzger, S.; Mahajan, R.; Melchior, F.; Jaenicke, R.; Becker, J., Structure determination of the small ubiquitin-related modifier SUMO-1. J Mol Biol 1998, 280 (2), 275–286.

28. Mossessova, E.; Lima, C. D., Ulp1-SUMO crystal structure and genetic analysis reveal conserved interactions and a regulatory element essential for cell growth in yeast. Molecular Cell 2000, 5 (5), 865–876.

29. Thomas, W. E.; Trintchina, E.; Forero, M.; Vogel, V.; Sokurenko, E. V., Bacterial adhesion to target cells enhanced by shear force. Cell 2002, 109 (7), 913–923.

30. Kim, J.; Zhang, C.-Z.; Zhang, X.; Springer, T. A., A mechanically stabilized receptor–ligand flex-bond important in the vasculature. Nature 2010, 466 (7309), 992.

31. Dorr, B. M.; Ham, H. O.; An, C.; Chaikof, E. L.; Liu, D. R., Reprogramming the specificity of sortase enzymes. Proc Natl Acad Sci U S A 2014, 111 (37), 13343–8.

32. Parks, T. D.; Howard, E. D.; Wolpert, T. J.; Arp, D. J.; Dougherty, W. G., Expression and purification of a recombinant tobacco etch virus Nla proteinase: biochemical analyses of the full-length and a naturally occurring truncated proteinase form. Virology 1995, 210, 194–194.

33. Milla, M. E.; Leesnitzer, M. A.; Moss, M. L.; Clay, W. C.; Carter, H. L.; Miller, A. B.; Su, J.-L.; Lambert, M. H.; Willard, D. H.; Sheeley, D. M., Specific sequence elements are required for the expression of functional tumor necrosis factor-α-converting enzyme (TACE). Journal of Biological Chemistry 1999, 274 (43), 30563–30570.

34. Tanner, N. A.; van Oijen, A. M., Visualizing DNA Replication at the Single-Molecule Level. In Single Molecule Tools, Part B:Super-Resolution, Particle Tracking, Multiparameter, and Force Based Methods, 2010; pp 259–278.

35. Chandradoss, S. D.; Haagsma, A. C.; Lee, Y. K.; Hwang, J. H.; Nam, J. M.; Joo, C., Surface passivation for single-molecule protein studies. J Vis Exp 2014, (86).

36. Gambino, S.; Mousley, B.; Cathcart, L.; Winship, J.; Loparo, J. J.; Price, A. C., A single molecule assay for measuring site-specific DNA cleavage. Anal Biochem 2016, 495, 3–5.

37. Hermanson, G. T., Bioconjugate Techniques. 3 ed.; Academic Press: 2013.

38. David Row, R.; Roark, T. J.; Philip, M. C.; Perkins, L. L.; Antos, J. M., Enhancing the efficiency of sortase-mediated ligations through nickel-peptide complex formation. Chem Commun (Camb) 2015, 51 (63), 12548–51.

39. Levary, D. A.; Parthasarathy, R.; Boder, E. T.; Ackerman, M. E., Protein-protein fusion catalyzed by sortase A. PLoS ONE 2011.

40. Kim, H.; Loparo, J. J., Multistep assembly of DNA condensation clusters by SMC. Nat Commun 2016, 7, 10200.

41. Edelstein, A. D.; Tsuchida, M. A.; Amodaj, N.; Pinkard, H.; Vale, R. D.; Stuurman, N., Advanced methods of microscope control using muManager software. J Biol Methods 2014, 1(2).

42. Strick, T. R.; Allemand, J. F.; Bensimon, D.; Bensimon, A.; Croquette, V., The elasticity of a single supercoiled DNA molecule. Science 1996, 271 (5257), 1835–1837.

43. Crocker, J. C.; Grier, D. G., Methods of digital video microscopy for colloidal studies. J Colloid Interface Sci 1996.

44. Pelletier, V.; Gal, N.; Fournier, P.; Kilfoil, M. L., Microrheology of microtubule solutions and actin-microtubule composite networks. Phys Rev Lett 2009, 102 (18), 188303.

45. Tarantino, N.; Tinevez, J. Y.; Crowell, E. F.; Boisson, B.; Henriques, R.; Mhlanga, M.; Agou, F.; Israel, A.; Laplantine, E., TNF and IL-1 exhibit distinct ubiquitin requirements for inducing NEMO-IKK supramolecular structures. J Cell Biol 2014, 204 (2), 231–45.

